# Neurite density but not myelination of specific fiber tracts links polygenic scores to general intelligence

**DOI:** 10.1101/2025.01.15.633108

**Authors:** Christina Stammen, Javier Schneider Penate, Dorothea Metzen, Maurice J. Hönscher, Christoph Fraenz, Caroline Schlüter, Onur Güntürkün, Robert Kumsta, Erhan Genç

## Abstract

White matter is fundamental for efficient and accurate information transfer throughout the human brain and thus crucial for intelligence. Previous studies often demonstrated associations between fractional anisotropy (FA) as a metric of white matter “microstructural integrity” and intelligence, but it is still unclear, whether this relation is due to greater axon density, parallel, homogenous fiber orientation distributions, or greater myelination since all of these measures influence FA. Using neurite orientation dispersion and density imaging (NODDI) and myelin water fraction (MWF) imaging data, we analyzed the microstructural architecture of intelligence in more detail in a sample of 500 healthy young adults. Furthermore, we were interested whether specific white matter microstructural indices play intermediary roles in the pathway that links genetic disposition for intelligence to phenotype. Thus, we conducted for the first time mediation analyses investigating whether neurite density (NDI), orientation dispersion (ODI), and MWF of 64 white matter fiber tracts mediate the effects of polygenic scores for intelligence (PGS_GI_) on general intelligence. By doing so, we showed that NDI, but not ODI or MWF of white matter fiber tracts was significantly associated with general intelligence and that the NDI of six fiber tracts mediated the relation between genetic variability and *g*. These findings are a crucial step forward in decoding the neurogenetic underpinnings of general intelligence, as they identify that neurite density of specific fiber tracts relates polygenic variation to *g*, whereas orientation dispersion and myelination did not.

## Introduction

Intelligence usually refers to the “[…] ability to understand complex ideas, to adapt effectively to the environment, to learn from experience, to engage in various forms of reasoning, to overcome obstacles by taking thought” [1]. It is one of the most studied human phenotypes and the last decades have produced cognitive tests in abundance to measure intelligence [2]. Although the various tests capture different aspects of intelligence, such as reasoning abilities or processing speed [3], individuals who perform well on one test tend to achieve high scores on other cognitive tests, regardless of the skills required [4,2]. Spearman [4] concluded that there was a factor of general intelligence, *g*, positioned at the apex of the hierarchy, with broader cognitive domains situated below and more specific cognitive abilities at the base [3,5]. The *g* factor illustrates the generalist character of intelligence [6], which is also reflected in the numerous associated life outcomes, such as school performance [7], job performance [8], socioeconomic success [9], income [10], or physical health [11]. In addition to the predictive value of intelligence, interindividual differences have been shown to be stable across the lifespan [12]. Consequently, there have been countless research efforts to uncover the neurogenetic mechanisms from which interindividual differences in intelligence arise.

A well-known framework for understanding intelligence on a neurobiological level provides the Parieto-Frontal Integration Theory of intelligence (P-FIT) [13]. This model suggests that intelligence is based on numerous areas throughout the cortex, especially in frontal and parietal regions, which are strongly interconnected and enable efficient information transfer. The idea that a brain network underlies intelligence also emphasized the role of structural connections, which are present in the form of white matter fiber tracts [13].

One technique to quantify white matter properties is diffusion-weighted imaging (DWI), whose advent has ushered in a new era of white matter brain imaging studies [2,14]. As the diffusion of water molecules is a three-dimensional process that includes random translational motion of molecules in space, diffusion anisotropy allows inferences about the presence of obstacles such as axons and their myelin sheaths [14]. Most studies use fractional anisotropy (FA) as a summative metric to describe white matter microstructural integrity [15]. In this context, higher FA values suggest more parallel diffusion pathways [16,17].

The relation between FA and intelligence has been widely studied [18–35]. The majority of studies reported positive associations between average FA values from many major white matter fiber tracts and cognitive performance [15]. Cox et al. [26], for example, used a large sample from the UK Biobank study and reported significant positive associations between FA and general intelligence in 25 out of 27 white matter fiber tracts. As summarized by Genç, Fraenz [15], FA values of the genu and the splenium of the corpus callosum, the uncinate fasciculus, and the superior longitudinal fasciculus were most often associated with differences in intelligence. In one recent study, Stammen et al. [34] showed robust associations between FA and general intelligence in four independent samples, located around the left-hemispheric forceps minor, superior longitudinal fasciculus, and cingulum-cingulate gyrus.

Since general intelligence appears to be robustly associated with higher FA values and thus stronger anisotropic diffusion patterns, Stammen et al. [34] formulated three possible explanations as to why higher FA values could be associated with higher intelligence. First, the relation could be due to greater axon density, enabling more parallel information processing by providing more pathways to think through problems relatively simultaneously. Second, the relation could be due to parallel, homogenous fiber orientation distributions that run directly from one brain region to another, thereby enabling more direct and efficient information transfer throughout the brain. Third, the relation could be due to greater myelination, enabling faster information processing speed and signal conduction velocity [36,37]. However, the exact neurobiological basis causing FA signal differences remains unclear, so the hypotheses put forward cannot be tested by looking at FA values alone. FA is a multifaceted and non-specific metric influenced by various physiological factors, including axon diameter, fiber density, myelin concentration, or the distribution of fiber orientation [38,14,39,40].

Recent advances in neuroimaging offer promising techniques that allow more differentiated and specific conclusions to be drawn about the microstructure of white matter and the three hypotheses mentioned [40]. Neurite orientation dispersion and density imaging (NODDI) [41] and myelin water fraction (MWF) imaging [42,37] both are non-invasive, but take advantage of the unique patterns that water molecules create in different environments.

NODDI utilizes a three-compartment model that differentiates between intra-neurite, extra-neurite, and cerebrospinal fluid (CSF) environments based on a multi-shell high-angular-resolution diffusion imaging protocol [41]. While isotropic diffusion occurs mainly in regions with CSF, intracellular compartments are characterized by stick-like or cylindrically symmetrical diffusion, as the water molecules are restricted by the membranes of neurites, and extracellular compartments by hindered diffusion, as there are many cellular membranes of somas and glia cells. The different diffusion properties allow the estimation of NODDI markers as approximations for different aspects of neurite morphology, such as the neurite density index (NDI), which represents the volume fraction of intra-neurite environments, and the orientation dispersion index (ODI), which quantifies the angular variation of neurite dispersion [41]. Histological studies supported the validity of the NODDI model [43].

MWF imaging relies on signals from myelin water, as signals from larger molecules such as lipids and proteins (main components of myelin) quickly decay to zero [42,37]. Unmyelinated neurons and glia cells have single bilayer membranes, whereas myelinated neurons possess multi-bilayer membranes, with roughly 40% of their mass consisting of compartmentalized water [42]. Although there is currently no method capable to measure the myelin bilayer directly, MWF imaging is considered the method of choice to estimate myelin content *in vivo* [42,37]. Studies validated the specificity of MWF imaging mapping the myelin bilayer by comparison with histological results [44–47] and demonstrated that MWF imaging has good reliability [48–51].

Thus, the question of whether the association between higher FA values and higher intelligence is due to greater axon density, parallel, homogenous fiber orientation distributions, or greater myelination can be answered by associating general intelligence with the corresponding indices NDI, ODI, and MWF. However, there are few to no studies that have used NODDI or MWF imaging in relation to intelligence.

Regarding NODDI, Genç et al. [52] were the first to analyze the microstructural architecture of intelligence. They showed that higher intelligence was associated with lower NDI and ODI in the gray matter and thus concluded that the neuronal circuitry linked to higher intelligence is structured in a sparse and efficient way [52]. For white matter, which is more relevant for this paper, associations between NODDI metrics and individual cognitive functions such as paired associate learning [53], episodic memory [54], or processing speed [54] have been reported in healthy adults. Callow et al. [55] found that the ODI of the cerebellar peduncle was significantly associated with fluid, but not crystallized cognition in healthy young adults.

To the best of our knowledge, there is no paper to date that analyzed MWF with regard to general intelligence in healthy adults. Penke et al. [20] were the first to use a biomarker of myelin, magnetization transfer ratio (MTR), to study intelligence in a sample of healthy older people. They were able to show that MTR correlated significantly with general intelligence. However, although a change in myelin content causes a change in MTR, MTR is also influenced by other pathological factors [42,56,57], so MTR is not as specific to myelin as MWF [42,37]. The relation between MWF and cognitive abilities in healthy adults has only been studied for specific cognitive functions such as processing speed [58], executive functions [59], or memory performance [60], but not for general intelligence.

To better understand the microstructural architecture of general intelligence, our study aimed to analyze the relation between general intelligence and NDI, ODI, and MWF within the same sample, with all indices extracted from the same 64 white matter fiber tracts. To further complete the picture, we did not limit our analyses to the relation between the brain and intelligence, but included genes and directly considered the triad between genes, the brain, and behavior via mediation analyses.

General intelligence is a highly heritable trait, with inherited differences in desoxyribonucleic acid (DNA) sequence explaining for about half of the variance in intelligence across all ages [61,6,62,2]. However, heritability of intelligence is not due to few individual genes, but results from thousands of genetic variants, mostly single nucleotide polymorphisms (SNPs), whose small effects on the variation of intelligence add up [2,6]. Polygenic scores (PGS) offer the possibility to account for this highly polygenic architecture by aggregating the effects of different SNPs across the genome into a summarized measure [63]. Results of genome-wide association studies (GWAS), used to identify which SNPs throughout the genome are statistically associated with a particular trait, show which of the two alleles for a SNP is positively associated with the trait (called increasing allele) and provide effect sizes for each SNP [2,6]. A PGS is constructed by summing the number of increasing alleles associated with intelligence across SNPs and weighting them by the respective effect size obtained from GWAS [2,6]. PGS for intelligence, derived from one of the largest GWAS to date based on 269,867 individuals, explain up to 5.2% of variance in general intelligence in independent samples [64].

Genetic correlations show that the genetic variants associated with intelligence are partially consistent with those associated with brain structure [2,65]. White matter microstructure has been shown to be highly polygenic as well and positive genetic correlations with higher intelligence have been identified [2,66–68]. The gene sets significantly associated with intelligence include neurogenesis, neuron differentiation, central nervous system neuron differentiation, regulation of nervous system development, positive regulation of nervous system development, and regulation of synapse structure or activity [64]. While other GWAS analyses on intelligence reported similar gene sets [65,69], gene sets such as myelin sheath or regulation of myelination were not found to be significantly related to intelligence in any study. Lee et al. [70], who performed a large GWAS on educational attainment, also found no gene sets related to glial cells that were positively enriched and concluded that differences in cognition may not necessarily be driven by differences in myelination and thus transmission speed.

Mediation analyses blaze the best trail to investigate the relation between PGS for general intelligence (PGS_GI_), microstructural white matter indices (NDI, ODI, and MWF), and general intelligence. However, there are only few mediation studies in healthy adults investigating the relation between genes, the brain, and intelligence and those that exist have focused their analyses on mediation effects of total brain volume, surface area, cortical thickness, white matter fiber network efficiency, and functional efficiency [71–75]. Although Genç et al. [72] found the white matter fiber network efficiency of two brain areas to be mediators regarding the effects of PGS on general intelligence using data from the same sample, they did not include white matter microstructure indices in their analyses, and to the best of our knowledge, other studies have not used such indices in mediation analyses in healthy adults either.

To summarize, NODDI and MWF imaging, with their indices NDI, ODI, and MWF, offer opportunities to analyze the microstructural architecture of intelligence on a new level and draw more differentiated and specific conclusions regarding the question whether the association between higher FA values and higher intelligence is due to greater axon density (NDI), parallel, homogenous fiber orientation distributions (ODI), or greater myelination (MWF). While these opportunities have only been used selectively for NODDI, to our knowledge there is no study that has investigated the relation between MWF and general intelligence in healthy young adults, so we performed this analysis here for the first time. Furthermore, we expanded the existing literature on mediation analyses regarding the relations between genes, the brain, and intelligence to include white matter microstructure. We analyzed the effects of PGS_GI_ on general intelligence and tested the mediating role of NDI, ODI, and MWF of 64 white matter fiber tracts in a large sample of at least 500 individuals. Thus, this study presents the first mediation analyses that give insight whether white matter microstructure indices provide a biological pathway through which our genetics influence general intelligence.

## Methods

### Participants

The sample consisted of 557 adults with a mean age of 27.33 years (SD = 9.43 years; range: 18-75 years, 503 right-handers), including 283 men (mean age = 27.71 years; SD = 9.86 years, 246 right-handers) and 274 women (mean age = 26.94 years; SD = 8.96 years, 257 right-handers). It has previously been used to investigate relations between genetic variability, brain properties, and intelligence [72]. Handedness was assessed using the Edinburgh Handedness Inventory [76]. Participants were mostly university students of different majors (mean years of education = 17.14 years; SD = 3.12 years), who were either financially compensated for their participation or received course credits. Individuals who had insufficient German language skills or reported having done any of the employed intelligence tests within the last five years were excluded from the study. Health status was assessed by a self-report questionnaire. Individuals were also not admitted to the study if they or any of their close relatives suffered currently or in the past from neurological and/or mental illnesses. The study protocol was approved by the Local Ethics Committee of the Faculty of Psychology at Ruhr University Bochum (vote Nr. 165). All participants gave written informed consent and were treated according to the Declaration of Helsinki.

### Acquisition and analysis of behavioral data

General intelligence was assessed by the use of four paper-and-pencil tests. The tests were conducted in a quiet and well-lit room.

#### I-S-T 2000 R

The Intelligenz-Struktur-Test 2000 R (I-S-T 2000 R) [77] is a well-established German intelligence test battery, requiring about 2.5 hours to complete. It evaluates various aspects of intelligence as well as general intelligence and is largely comparable to the internationally established Wechsler Adult Intelligence Scale Forth Edition [78]. The majority of included cognitive test items are presented in multiple-choice format. The test consists of a basic and an extension module. Within the basic module, verbal, numerical, and figural abilities are each assessed by three different mental reasoning tasks of 20 items. Verbal intelligence is assessed by tasks in which participants must complete sentences (IST_SEN), find analogies (IST_ANA), and recognize similarities (IST_SIM). Numerical intelligence is assessed by tasks involving arithmetic calculations (IST_CAL), number series (IST_SER), and mathematical equations to which arithmetic signs need to be added (IST_SIG). Figural intelligence is assessed by tasks in which participants must select and reassemble parts of a cut-up figure (IST_SEL), mentally rotate and match three-dimensional objects (IST_CUB), and solve matrix-reasoning problems (IST_MAT). In addition, retention (IST_RET) is assessed by ten verbal and 13 figural items. Here, participants must memorize series of words or figure pairs. The extension module comprises 84 multiple-choice questions on six knowledge facets (art/literature, economy, geography/history, mathematics, science, and daily life) and measures general knowledge (IST_KNO). Reliability estimates (Cronbach’s α) are between .88 and .96 for subtests and composite scores. The recent norming sample consists of about 5800 individuals for the basic module and 661 individuals for the extension module. The age range in the norming sample is between 15 and 60 years and both sexes are represented equally [77].

#### BOMAT-Advanced Short

The Bochumer Matrizentest (BOMAT) [79] is a non-verbal German intelligence test which is widely used in neuroscientific research [79–82,52,83,84] and whose structure resembles that of the well-established Raven’s Advanced Progressive Matrices [85]. Within the framework of our study, we employed the advanced short version, which is characterized by high discriminatory power in samples with generally high intellectual abilities, thus avoiding possible ceiling effects [52,83]. The test comprises two parallel forms with 29 matrix-reasoning items. Each item shows a 5-by-3 matrix composed of elements arranged according to a specific but unspecified rule. One field within the matrix is empty and needs to be filled with one of six provided elements that follows the rule. The participants were assigned to one of the two parallel forms and had to complete as many matrices as possible within a time limit of 45 minutes. Split-half reliability of the BOMAT is .89, Cronbach’s α is .92, and reliability between the parallel forms is .86. The recent norming sample consists of about 2100 individuals with an age range between 18 and 60 years and equal sex representation [79].

#### BOWIT

The Bochumer Wissenstest (BOWIT) [86] is a German general knowledge questionnaire. It assesses eleven different knowledge facets, from two major domains. The four facets biology/chemistry, mathematics/physics, nutrition/exercise/health, and technology/electronics are assigned to the scientific-technical knowledge domain. The social and humanistic knowledge domain includes seven facets: arts/architecture, civics/politics, economies/laws, geography/logistics, history/archaeology, language/literature, and philosophy/religion. The BOWIT is available in two parallel test forms, in which each knowledge facet is represented by 14 multiple-choice questions. To measure general knowledge as precisely as possible, all participants had to complete both test forms, resulting in 308 items. The BOWIT shows reliability estimates greater than .90: split-half reliability is reported as .96, Cronbach’s α .95, test-retest reliability .96, and parallel-form reliability .91. The recent norming sample consists of about 2300 individuals with an age range between 18 and 66 years and equal sex representation [86].

#### ZVT

The Zahlenverbindungstest (ZVT) [87] is a trail-making test used to assess the cognitive processing speed of both children and adults. After completing two short sample tasks, four main tasks are assessed. Here, participants connect numbers from 1 to 90 according to a specific rule as fast as possible. The processing times for the four tasks are averaged to obtain an overall measure of processing speed. The reliability across the four tasks is reported as .95 in adults. The six-month retest-reliability is reported to be between .84 and .90. The recent norming sample consists of about 2109 individuals with an age range between eight and 60 years and equal sex representation [87].

#### Computation of the general intelligence factor, *g*

We computed the general intelligence factor to provide a comparable and robust measure of intelligence. When included tests measure intelligence broadly enough, *g* factors derived from different test batteries are statistically equivalent [88,89]. As described in Stammen et al. [34], we used the intelligence test scores to compute *g* factor scores for every participant. After regressing age, sex, age*sex, age^2^, and age^2^*sex from the test scores, we used the standardized residuals to develop a hierarchical factor model via exploratory factor analysis. Subsequently, we performed confirmatory factor analysis to assess model fit using the chi-square (Х^2^) statistic as well as the fit indices Root Mean Square Error of Approximation (RMSEA), Standardized Root Mean Square Residual (SRMR), Comparative Fit Index (CFI), and Tucker-Lewis Index (TLI). The evaluation of model fit yielded good fit. Although the chi-square (Х^2^) statistic assessing the magnitude of discrepancy between the model-implied variance-covariance matrix and the empirically observed variance-covariance matrix [90] was significant (Х^2^(64) = 127.97, *p* < .001), this did not itself show poor model fit since the chi-square (Х^2^) statistic is a direct function of sample size meaning that the probability of rejecting any model increases with greater sample size [91,92]. The other fit indices (RMSEA = .042, SRMR = .033, CFI = .979, and TLI = .969) were all acceptable as values of RMSEA and SRMR less than .05 and values of CFI and TLI greater than .97 are considered good [90]. Based on the postulated confirmatory factor model shown in Figure 1, we calculated regression-based *g*-factor scores for every participant, winsorizing outliers [93].

**Fig. 1.**
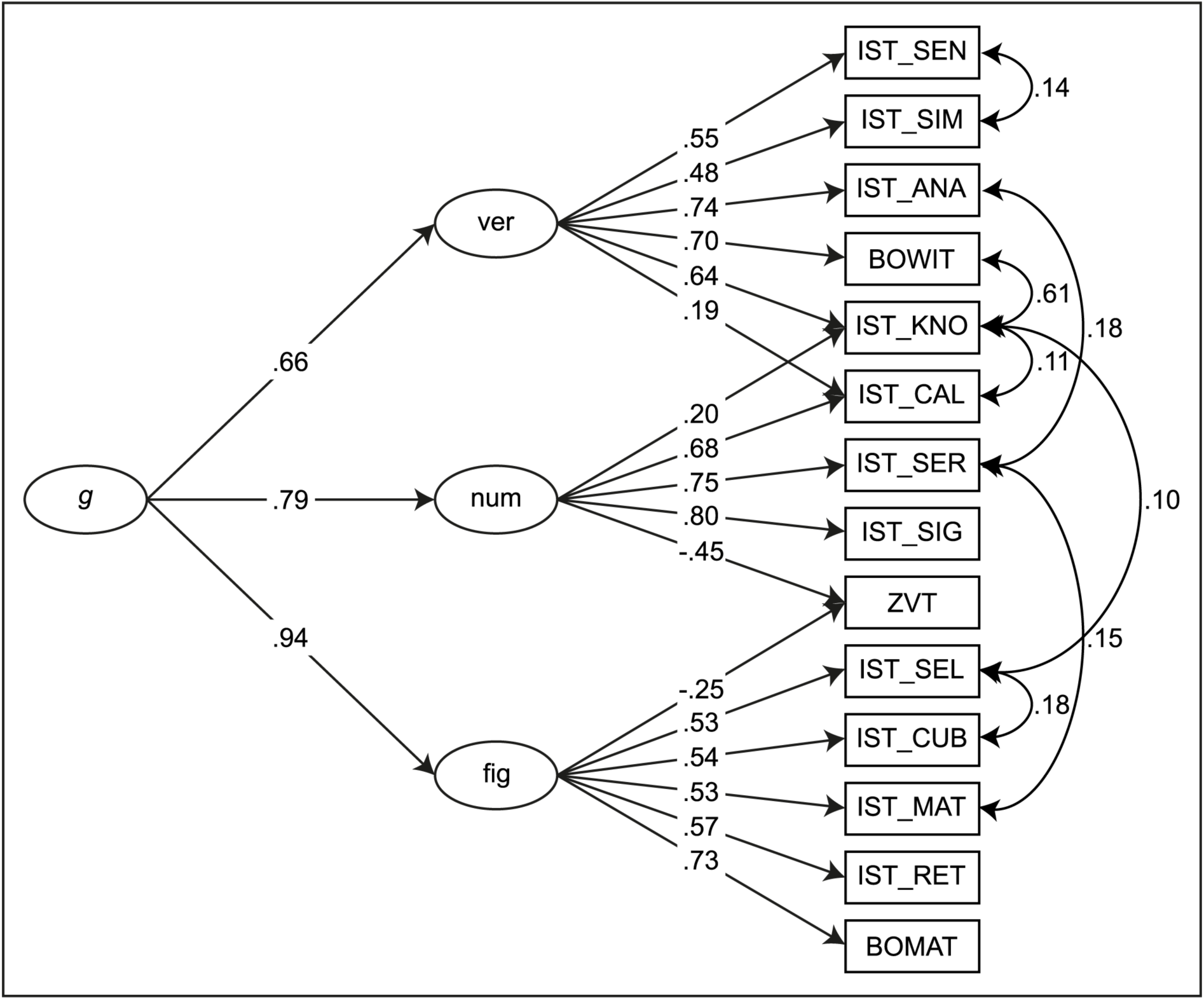
Confirmatory factor analytic model. *g* = general factor of intelligence, ver = verbal intelligence as broad cognitive domain, num = numerical intelligence as broad cognitive domain, fig = figural intelligence as broad cognitive domain, IST_SEN = subtest Sentence Completion of the I-S-T 2000 R, IST_SIM = subtest Similarities of the I-S-T 2000 R, IST_ANA = subtest Analogies of the I-S-T 2000 R, BOWIT = Bochumer Wissenstest, IST_KNO = parameter Knowledge of the I-S-T 2000 R, IST_CAL = subtest Calculations of the I-S-T 2000 R, IST_SER = subtest Number Series of the I-S-T 2000 R, IST_SIG = subtest Numerical Signs of the I-S-T 2000 R, ZVT = Zahlenverbindungstest, IST_SEL = subtest Figure Selection of the I-S-T 2000 R, IST_CUB = subtest Cubes of the I-S-T 2000 R, IST_MAT = subtest Matrices of the I-S-T 2000 R, IST_RET = parameter Retentiveness of the I-S-T 2000 R, BOMAT = Bochumer Matrizentest

#### Intelligence level

Unfortunately, it is not possible to link *g* to the intelligence quotient (IQ) scale. Nevertheless, we used norming data of some tests to estimate the intelligence level. Norming data of the subtests of the I-S-T 2000 R revealed that the sample’s mean IQ was 115 (SD = 13.0). The fact that our sample had a mean score one standard deviation above average may have impacted the associations with the polygenic scores and the white matter fiber tracts’ microstructure.

### DNA sampling and genotyping

We used exfoliated cells that were brushed from the participants’ oral mucosa for genotyping. DNA isolation was done with QIAamp DNA mini Kit (Qiagen GmbH, Hilden, Germany). Genotyping was executed with the Illumina Infinium Global Screening Array 1.0 with MDD and Psych content (Illumina, San Diego, CA, USA) at the Life & Brain facilities (Bonn, Germany) and yielded 745,747 SNPs. Filtering was conducted with PLINK 1.9 [94,95] by eliminating all SNPs with a minor allele frequency (MAF) of < 0.01, deviating from Hardy-Weinberg equilibrium (HWE) with a *p*-value of < 1*10^-6^, and missing data > 0.02. Participants were removed with > 0.02 missingness, sex-mismatch, and heterozygosity rate > |0.2|. A high quality (HWE *p* > 0.02, MAF > 0.2, missingness = 0) and linkage disequilibrium (LD) pruned (*r*^2^ = 0.1) SNP set was used for filtering for relatedness and population structure. In pairs of related subjects, pi hat > 0.2 was used to exclude subjects randomly. Principal components (PCs) were generated to control for population stratification. Participants who deviated more than |6| standard deviations from the mean on at least one of the first 20 PCs were classified as outliers and excluded. The final data set consisted of 519 participants and 498,760 SNPs. The samples’ filtered genotype data was submitted for imputation to the Michigan Imputation server [96] using the European population of the Haplotypes Reference Consortium panel (r1.1 2016; hg19) and a R^2^ filter of 0.3. We chose Eagle 2.4 for phasing and Minimac4 for imputation. After a final MAF < 0.01 filtering step, 5,338,876 SNPs were available for analysis.

### Polygenic scores

We generated genome-wide PGS for each participant using publicly available summary statistics for general intelligence [N = 269,867; 64]. General intelligence PGS (PGS_GI_) were computed as weighted sums of each subject’s trait-associated alleles across all SNPs using PRSice 2.1.6 [97]. Specifically, we computed best-fit PGS_GI_ that showed the strongest association with general intelligence [72,84]. We applied a *p*-value threshold (PT) for the inclusion of SNPs that was chosen empirically by carrying out multiple linear regression analyses iteratively for the range of PT 5*10^-8^ to 0.5 in steps of 5*10^-5^. The predictive power of the PGS_GI_ was assessed by the “incremental *R*^2^” statistic [70]. Incremental *R*^2^ indicates the increase in the coefficient of determination (*R*^2^) when the PGS_GI_ is added as a covariate to a regression model predicting general intelligence together with a number of baseline control variables (here: age, sex, and the first four PCs of population stratification). Linear parametric methods were chosen for all statistical analyses in PRSice. Testing was two-tailed with an α-level of *p* < .05. The PGS_GI_ with the greatest predictive power, explaining the maximum amount of *g* variance, was chosen for further analyses. The best-fit threshold selected for PGS_GI_ was 0.0062, which resulted in 4,659 SNPs being included in the calculation of PGS_GI_. PGS_GI_ explained 4.37% of variance of general intelligence in our sample (*p* < .001). A distribution of the PGS_GI_ is shown in Figure S1.

### Acquisition and analysis of imaging data

All images were collected within one session on a Philips 3T Achieva scanner at the Bergmannsheil Hospital in Bochum, Germany, using a 32-channel head coil.

#### Multi-shell diffusion-weighted imaging

For the analysis of NODDI coefficients, a diffusion-weighted three-shell image was acquired using echo-planar imaging (EPI) with the following parameters: time repetition (TR) = 7652 ms; time echo (TE) = 87 ms; flip angle = 90°; 60 slices; matrix size = 112 x 112; voxel size = 2 x 2 x 2 mm; parallel imaging sensitivity encoding (SENSE) factor = 2; direction of acquisition = anterior-posterior (AP). Diffusion weighting was uniformly distributed along 120 directions (20 directions with a *b*-value of 1000 s/mm^2^; 40 directions with a *b*-value of 1800 s/mm^2^; 60 directions with a *b*-value of 2500 s/mm^2^). We used the multiple acquisitions for standardization of structural imaging validation and evaluation toolbox (MASSIVE toolbox) [98] to generate all diffusion directions within and between shells non-collinear to each other. Additionally, eight volumes with no diffusion weighting (*b*-value of 0 s/mm^2^) were acquired for the purpose of motion correction and computation of NODDI coefficients. Diffusion-weighted data was collected with reversed phase-encode directions, resulting in pairs of images with distortions going in opposite directions. Total acquisition time was 18 minutes.

Diffusion-weighted data were prepared for NODDI coefficients via a preprocessing pipeline comprising the following steps. First, images were corrected for signal drift [99] using ExploreDTI [100]. Second, we utilized the topup command from the Oxford Centre for Functional Magnetic Resonance Imaging of the Brain’s (FMRIB’s) Software Library (FSL) toolbox (version 6.0.7.7; https://fsl.fmrib.ox.ac.uk/fsl/docs/#/) [101,102] to estimate the susceptibility-induced off-resonance field based on pairs of images with opposite phase-encode directions. Third, the topup output was used in combination with the eddy command [103], which is also part of the FSL toolbox [101], to correct for susceptibility, eddy currents, and head movement. Importantly, we also performed outlier detection during this step to identify slices where signal had been lost due to head movement during the diffusion encoding [104].

We used the Microstructure Diffusion Toolbox (MDT; https://github.com/robbert-harms/MDT) [105,106] to compute NODDI coefficients. The advantage of MDT in comparison to the original NODDI toolbox in MATLAB [41] is that MDT is utilized on Graphics Processing Unit (GPU) cores and thus dramatically reduces estimation time. It is even faster than the AMICO toolbox [107] we used in previous studies [40,52,108]. By default, MDT uses the Offset Gaussian likelihood model and the Powell conjugate-direction optimization routine [109] for maximum likelihood estimation [105]. Specifically, we employed the implemented, three-part NODDI model of Zhang et al. [41] that distinguishes between intra-neurite, extra-neurite, and CSF environments. The NODDI technique is based on a two-level approach. First, the proportion of free moving water within each voxel is analyzed based on the diffusion signal obtained by the multi-shell high-angular-resolution imaging protocol [110–112,41]. This proportion is called isotropic volume fraction and reflects the amount of isotropic diffusion with Gaussian properties that mainly characterizes regions with a focus on CSF. Second, the remaining portion of the diffusion signal is assigned to one of the complementary fractions, either intra- or extra-neurite environment [41,110,111]. The amount of intra-neurite environments is quantified as the intra-neurite volume fraction or NDI. The intra-cellular compartment represents the amount of stick-like or cylindrically symmetric diffusion that occurs when water molecules are confined by the membranes of neurites and resembles the proportion of axonal density in white matter as shown by comparison with light microscopy and electron microscopy in histological samples [110]. Extra-neurite environments in the white matter are usually full of various types of glia cells and therefore characterized by hindered diffusion [110,111,41]. A NODDI’s summary statistic is the neurite ODI that quantifies angular variation of neurite orientation [41]. ODI is a measure of tortuosity that couples the intra-neurite space and the extra-neurite space and thus leads to an alignment or dispersion of the axons in the white matter [112,41]. Examples of NDI and ODI coefficient maps from a representative individual are illustrated in Figure 2 (upper-left corner).

**Fig. 2.**
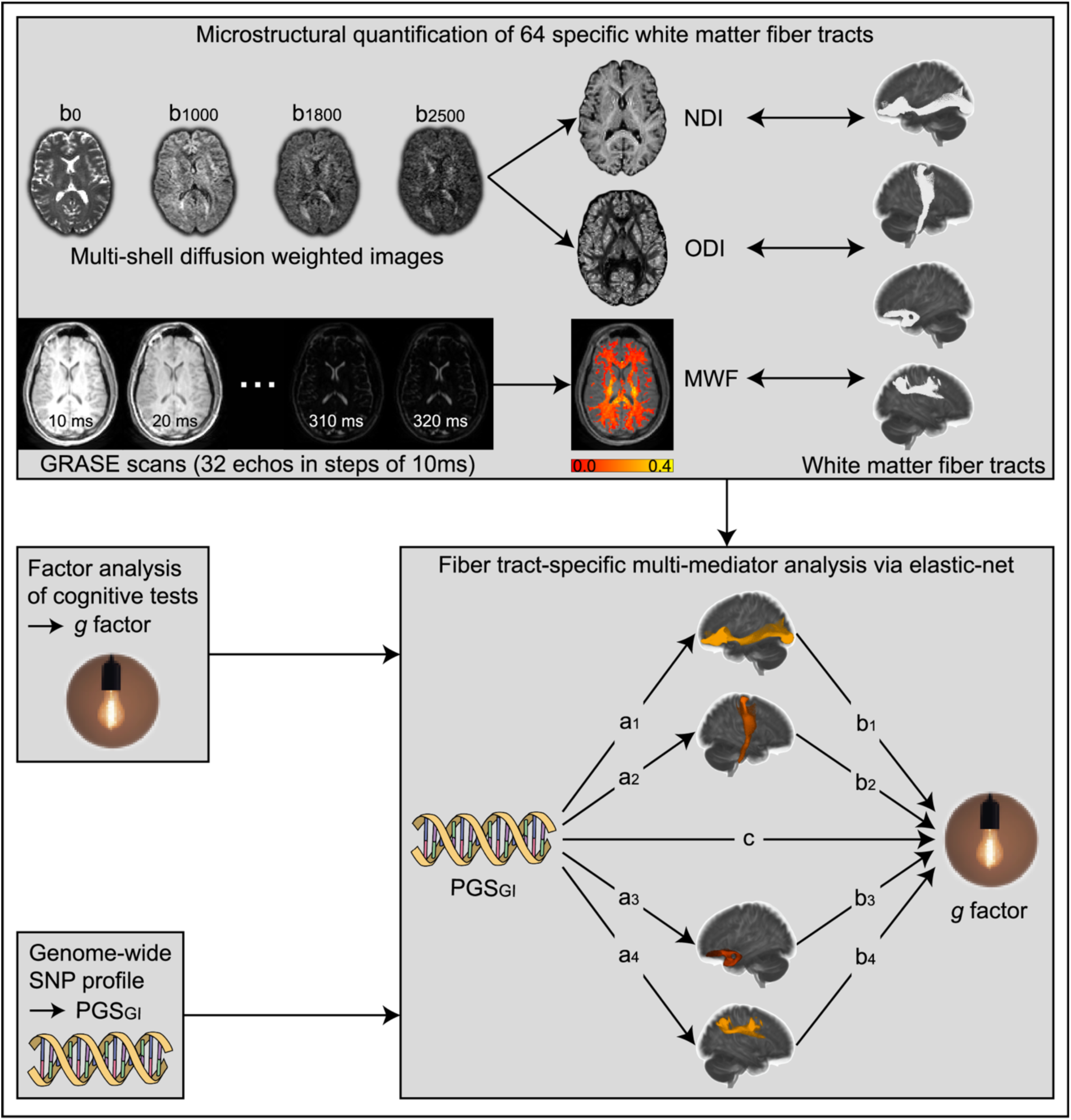
Processing steps of neuroimaging and statistical analyses. Multi-shell diffusion-weighted images were used to compute neurite density and neurite orientation dispersion indices (NDI and ODI). Images resulting from the three-dimensional (3D) multi-echo (ME) turbo gradient spin echo (GRASE) sequence were used to compute the myelin water fraction (MWF) for each voxel. 64 white matter fiber tracts provided by the population-based HCP-1065 probabilistic tract atlas [119,120] were downloaded, transformed in the respective native space, and served as anatomical references for the extraction of the microstructural properties NDI, ODI, and MWF. Fiber tract-specific multi-mediator analyses were performed via elastic-net regression for each microstructural property. General intelligence, quantified by the factor of general intelligence *g*, was the dependent variable, while the polygenic score of general intelligence (PGS_GI_) was the independent variable. NDI, ODI, or MWF values of each white matter fiber tract served as mediators

#### Myelin water imaging

A previously published 32 multi-echo (ME) three-dimensional (3D) turbo gradient spin echo (GRASE) sequence with refocusing angle sweep [113,112,40,114] was acquired with the following parameters: TR = 800 ms; TE = 32 echoes at 10 ms echo spacing (ranging from 10 to 320 ms); 60 slices; partial Fourier acquisition in both phase encoding directions; matrix size = 112 x 112; voxel size = 2 x 2 x 2 mm; parallel imaging SENSE factor = 2; direction of acquisition = right-left (RL). Total acquisition time for this sequence was eight minutes.

To assess the myelin content of the white matter fiber tracts, we used an in-house algorithm [114,40] written in MATLAB (version 8.5.0.197613 (R2015a), The MathWorks Inc., Natick, MA) to construct parameter maps representing the MWF for each voxel based on the 3D ME-GRASE sequence. As described in Prasloski et al. [113] in detail, this algorithm analyzed the ME decay curves voxel by voxel using multicomponent T2 analysis with simultaneous correction for contamination of the decay curves by stimulated echoes resulting from B1 inhomogeneity and imperfect refocusing pulses. Voxelwise ME decay curves were acquired from the 3D ME-GRASE images and transformed into a continuous T2-distribution by applying a regularized non-negative least squares (NNLS) approach [115,116]. We used an extended phase graph algorithm to take possible stimulated echoes due to non-ideal refocusing pulse flip angles into consideration [117,118,113]. During the fitting procedure, a regularization factor of 1.02 was applied to increase robustness of the ill-posed fitting problem and to assure smooth T2 amplitude distributions. T2 distributions were generated using 101 logarithmically spaced exponential decay base functions for echo decay with T2 values ranging from 0.01 to 2s. From the T2 distributions, the MWF was calculated for each voxel as the signal integral fraction between 10 and 40ms relative to the total T2 distribution integral (area under the curve). This resulted in whole-brain MWF maps for each subject, as exemplified in Figure 2 (upper-left corner).

### Quantification of microstructural properties in white matter fiber tracts

We used 64 white matter fiber tracts provided by the population-based HCP-1065 probabilistic tract atlas [119,120], from the official website (https://brain.labsolver.org/hcp_trk_atlas.html) as NIfTI files. This atlas displays for 64 different fiber tracts for each voxel the probability of being part of the respective white matter fiber tract compiled from the tractography of 1065 subjects [119], with underlying data (“1200 Subjects Data Release”) provided by the Human Connectome Project (HCP), WU-Minn Consortium (Principal Investigators: David van Essen and Kamil Ugurbil; 1U54MH091657), funded by the 16 United States National Institutes of Health (NIH) and Centers supporting the NIH Blueprint for Neuroscience Research and by the McDonnell Center for Systems Neuroscience at Washington University [121]. In a first step, fiber tracts’ NIfTI files were processed using DSI Studio (https://dsi-studio.labsolver.org) [119,120]. We resized the dimensions to match the International Consortium for Brain Mapping (ICBM) 2009a Nonlinear Asymmetric NIfTI template file [122]. The threshold for the fiber tracts’ probability was set at 0.50 to include only voxels that were part of major white matter tracts in at least half of the sample and exclude peripheral voxels that are more susceptible to intra- and intersubjective variability. We then binarized the fiber tracts and transformed them into a common space via FMRIB’s Linear Image Registration Tool (FLIRT) [123–125]. We chose the template MNI152_T1_1mm_brain within FSL, which is derived from 152 structural images that have been nonlinearly registered into the common Montreal Neurologic Institute (MNI) 152 standard space (1 x 1 x 1 mm). Starting from the MNI 152 standard space, we used FMRIB’s Nonlinear Image Registration Tool (FNIRT) [126] to nonlinearly transform the fiber tracts into the native space of the diffusion-weighted images as well as into the 3D-ME GRASE image space. Each participant’s aligned fiber tracts served as anatomical references from which NODDI coefficients and MWF coefficients were extracted (see Figure 2, upper-right corner).

### Statistical analyses

All statistical analyses were conducted in R Studio (version 2022.12.0.353) [127] with R version 4.2.2 (2022-10-31) [128]. The final data set included 501 participants (242 women; mean age = 27.30 years; SD = 9.22 years; 455 right-handers) as we only had usable genetic data from 519 subjects and analyzable MWF data from 539 subjects. Data points were treated as outliers if they deviated more than three interquartile ranges from the respective variable’s group mean (PGS_GI_, mean NDI of all 64 fiber tracts, mean ODI of all 64 fiber tracts, mean MWF of all 64 fiber tracts, *g*). In such cases, all data from the corresponding participant were removed from analysis. No subjects were excluded from analyses concerning PGS_GI_, NDI, MWF, and *g*, while one participant had to be excluded for analyses concerning PGS_GI_, ODI, and *g* (500 remaining subjects).

To investigate whether a set of specific white matter fiber tracts mediates the association between PGS_GI_ (independent variable) and general intelligence (dependent variable), we used exploratory mediation analysis by regularization (see Figure 2, lower-right corner), an approach developed to identify a set of mediators from a large pool of potential mediators without testing specific theory-based and predefined hypotheses [129,130]. Confirmatory theory-based approaches in general test models that have been specified in advance, rely on *p*-values to test statistical significance, require correction for multiple comparisons with respect to many possible mediators, and tend to overfit the data in the regression context, resulting in less generalizable solutions [130–132]. In contrast, exploratory mediation analysis by regularization is based on regularization and penalization techniques [130], such as the least absolute shrinkage and selection operator (lasso) [133]. It aims to improve the generalization ability of a model and prevent overfitting [134] by applying a penalty to effect sizes, resulting in small effect sizes being pushed to zero, leaving only strong non-zero effects.

A detailed explanation of this machine learning approach is provided by Serang et al. [130]. In short, this approach is based on a two-stage process. First, all potential mediators of interest are included in a multiple mediator model which is then fit using lasso resulting in the corresponding regression weights *a* and *b* being penalized [135]. The tuning parameter of the penalty term, lambda, is typically selected by testing a range of candidate values via *k*-fold cross-validation, an approach primarily utilized to prevent overfitting [130]. The data is divided into *k* different subsets. The model is then trained on *k*-1 subsets, while the *k*th subset is used as the testing set. This is repeated *k* times so that each subset serves as the testing set once. The value of lambda chosen is the one with the best fit resulting in the lowest prediction error. Since the mediation effect of a mediator corresponds to the product of the regression parameters *a* and *b*, the effect becomes zero if either the *a* or *b* parameter of a mediator is regularized to zero by the penalty. Those mediators with non-zero values of *a* and *b* after regularization, will be considered selected as mediators. While this approach makes it possible to eliminate mediators with small effect sizes, it also means that the effect sizes of the selected mediators are close to zero due to penalization and are therefore underestimated. To eliminate this potential bias, the second step is to refit the model using only the selected mediators without any penalization [130]. This allows unbiased estimates of effect sizes to be obtained [129].

Instead of using lasso regression, we employed elastic-net regression in our analyses. Elastic-net regression results from the combination of ridge regression [136] and lasso regression [133] and is thus another form of regularized regression [137] allowing better accuracy of prediction on future data and interpretation of the model due to parsimony in contrast to ordinary least squares estimates. While lasso regression can penalize a parameter to zero and is therefore suitable for models where many variables are assumed to have little or no effect on the dependent variable, ridge regression can only asymptotically shrink parameters towards zero and is therefore suitable for models where most variables are assumed to have a considerable effect on the dependent variable. Elastic-net regression is an useful approach when there are no clear, predefined hypotheses for all variables [137]. Unlike lasso regression, which tends to randomly select only one variable from a group of variables with high correlations between them, elastic-net regression outperforms lasso as it can select groups of correlated variables [137]. The latter was an important argument in our decision to use elastic-net regression instead of lasso regression.

We used the *xmed* function from the *regsem* package [138,129,130]. All variables were standardized and residualized for age, sex, age*sex, age^2^, age^2^*sex, and the first four PCs of population stratification. Age, sex, age^2^, and their interaction effects were used as control variables, as many studies have shown age- and sex-dependent changes in microstructural properties as well as myelination [139–144]. The first four PCs of population stratification were added to control the variability of the genetic origin of the sample [145]. We calculated three mediation models, where PGS_GI_ was always the independent variable, the NDI-, ODI-, or MWF-values of the 64 white matter fiber tracts each yielded the 64 mediators, and *g* was always the dependent variable. For all mediation models, the number of cross-validation subsets was set to *k* = 80, the threshold for detecting non-zero mediation effects was set to 0.001 (default), and the type of regression was set to elastic-net. All coefficients were re-estimated with *lavaan* [146] to avoid biased effect sizes.

In addition to the mediation effects, we were also interested in the direct effects of PGS_GI_ on the NODDI and MWF brain properties as well as the direct effects of the NODDI and MWF brain properties on intelligence (paths *a* and paths *b*). To identify variables with non-zero effects within path *a* and path *b* regressions, we followed a similar approach and set the threshold for detecting non-zero effects to 0.01 [72]. This threshold was chosen a little more liberally, since mediation effects are considerably smaller due to the multiplication of the regularized parameters *a* and *b* with values less than one. Again, all coefficients were re-estimated with *lavaan* [146].

## Results

### Neurite density index (NDI)

The results of the multiple mediator analysis via elastic-net showed that PGS_GI_ was associated with the NDI of 28 white matter fiber tracts (see Figure 3 and supplementary Table S1). All effects were positive, indicating that higher PGS_GI_ is associated with higher NDI (path *a*) and thus higher axonal packing density. Furthermore, the NDI of 18 white matter fiber tracts was linked to the *g* score (path *b*). The association was positive for 12 white matter fiber tracts and negative for the remaining six fiber tracts. A total of six white matter fiber tracts mediated the effects of PGS_GI_ on general intelligence (path *a*\**b*). While the five white matter fiber tracts middle longitudinal fasciculus (left hemisphere), cingulum parahippocampal parietal (right hemisphere), uncinate fasciculus (left hemisphere), cingulum parahippocampal parietal (left hemisphere), and superior longitudinal fasciculus three (right hemisphere) showed positive mediation effects, one white matter fiber tract, namely frontal aslant tract (left hemisphere), showed a negative mediation effect.

**Fig. 3.**
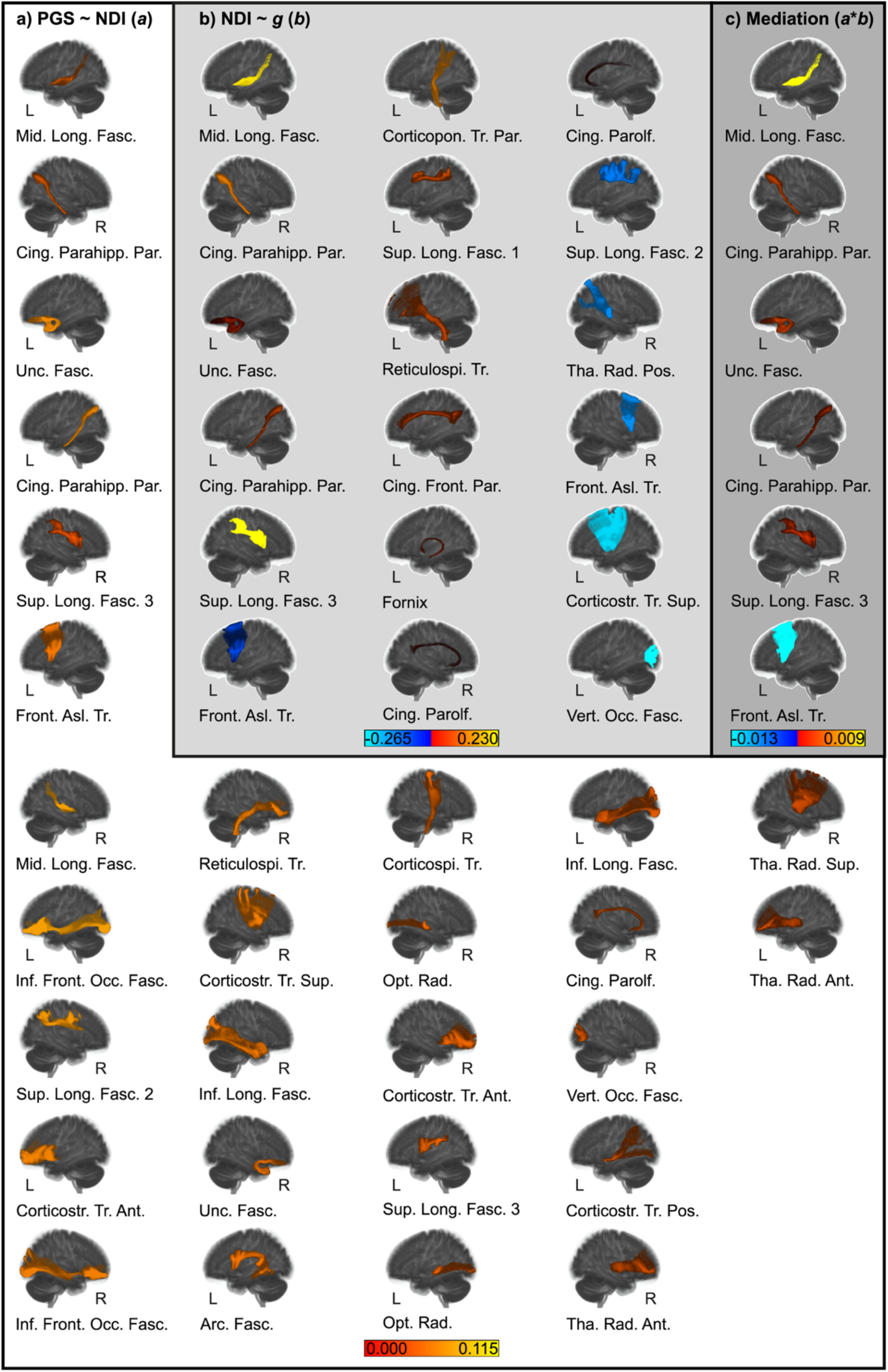
Results of the multiple mediator analysis via elastic-net with neurite density index (NDI) values of 64 white matter fiber tracts as mediators. The figure shows the results from path *a* analysis (block a) PGS ∼ NDI (*a*)) with white background, path *b* analysis (block b) NDI ∼ *g* (*b*)) with light gray background, and the mediation effects *a*\**b* (bock c) Mediation (*a*\**b*)) with gray background. White matter fiber tracts are shown in left (L) or right (R) sagittal view. Positive effects are depicted in red and yellow, negative effects are depicted in blue and light-blue. For a list of white matter fiber tracts and effect sizes see Table S1. Abbreviations: Arc. Fasc. = arcuate fasciculus; Cing. Front. Par. = cingulum frontal parietal; Cing. Parahipp. Par. = cingulum parahippocampal parietal; Cing. Parol. = cingulum parolfactory; Corticopon. Tr. Par. = corticopontine tract parietal; Corticospi. Tr. = corticospinal tract; Corticostr. Tr. Ant. = corticostriatal tract anterior; Corticostr. Tr. Pos. = corticostriatal tract posterior; Corticostr. Tr. Sup. = corticostriatal tract superior; Front. Asl. Tr. = frontal aslant tract; Inf. Front. Occ. Fasc. = inferior fronto-occipital fasciculus; Inf. Long. Fasc. = inferior longitudinal fasciculus; Mid. Long. Fasc. = middle longitudinal fasciculus; Opt. Rad. = optic radiation; Reticulospi. Tr. = reticulospinal tract; Sup. Long. Fasc. = superior longitudinal fasciculus; Tha. Rad. Ant. = thalamic radiation anterior; Tha. Rad. Pos. = thalamic radiation posterior; Tha. Rad. Sup. = thalamic radiation, superior; Unc. Fasc. = uncinate fasciculus; Vert. Occ. Fasc. = vertical occipital fasciculus

### Neurite orientation dispersion index (ODI)

For the ODI metric, the elastic-net analysis showed that there was an association between PGS_GI_ and the tracts’ ODI values in 16 white matter fiber tracts (see Figure 4a and supplementary Table S2). As for NDI, all effects were positive, indicating that higher PGS_GI_ is associated with higher ODI (path *a*). Higher ODI indicates a cytoarchitecture with highly dispersed neurites. The multiple mediator analysis revealed no white matter fiber tracts that showed a significant association between the tracts’ ODI values and general intelligence (path *b*). Consequently, no mediators (path *a*\**b*) for the association between PGS_GI_ and *g* could be identified.

**Fig. 4.**
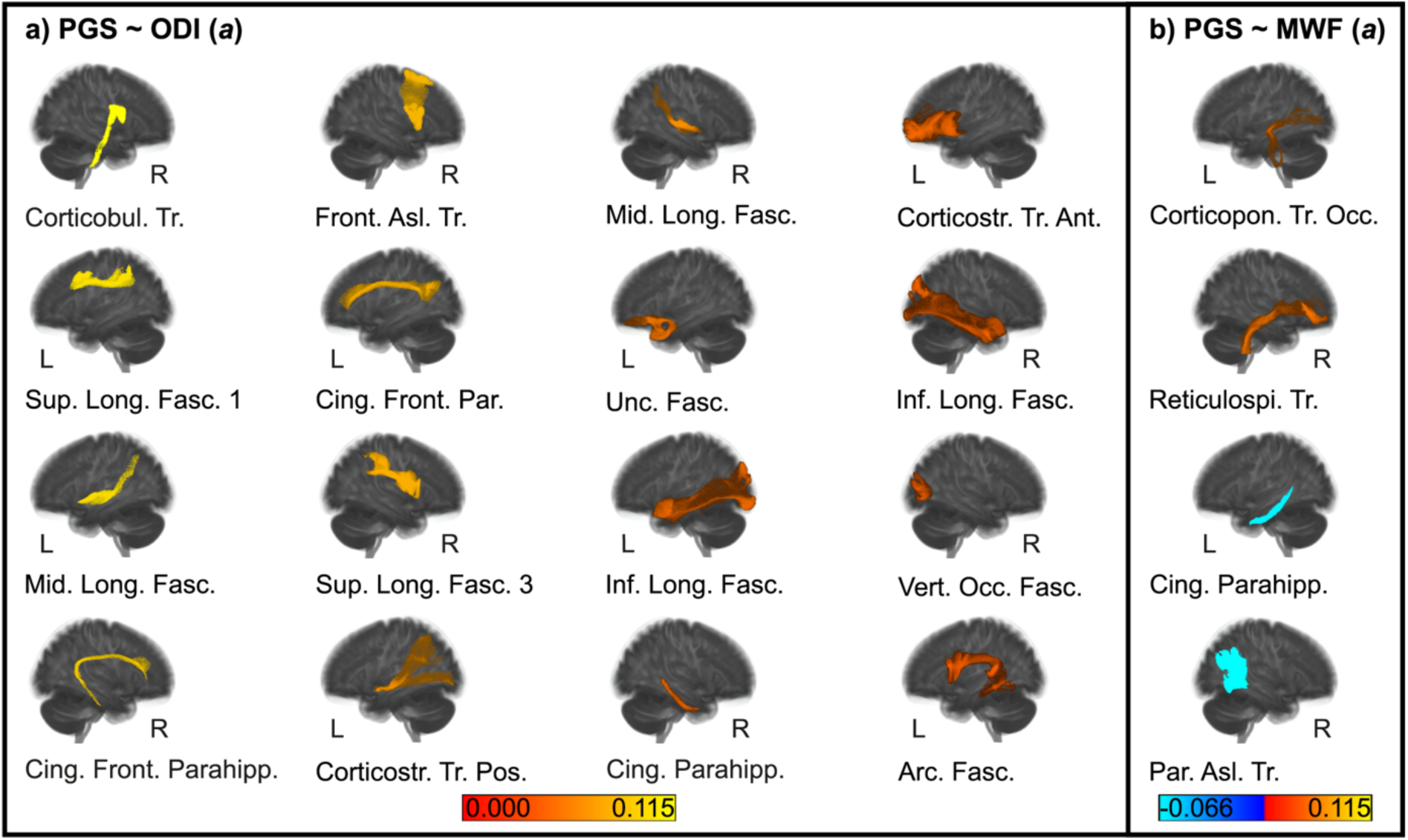
Results of the multiple mediator analysis via elastic-net with neurite orientation dispersion index (ODI) values (block a) PGS ∼ ODI (*a*)) or myelin water fraction (MWF) values (block b) PGS ∼ MWF (*a*)) of 64 white matter fiber tracts as mediators. The figure only shows the results from paths *a* analyses, as no significant associations could be identified for paths *b* and thus also not for paths *a*\**b*. White matter fiber tracts are shown in left (L) or right (R) sagittal view. Positive effects are depicted in red and yellow, negative effects are depicted in blue and light-blue. For a list of white matter fiber tracts and effect sizes see Table S2 for ODI and Table S3 for MWF. Abbreviations: Arc. Fasc. = arcuate fasciculus; Cing. Front. Parahipp. = cinglum frontal parahippocampal; Cing. Front. Par. = cingulum frontal parietal; Cing. Parahipp. = cingulum parahippocampal; Corticobul. Tr. = corticobulbar tract; Corticopon. Tr. Occ. = corticopontine tract occipital; Corticostr. Tr. Ant. = corticostriatal tract anterior; Corticostr. Tr. Pos. = corticostriatal tract posterior; Front. Asl. Tr. = frontal aslant tract; Inf. Long. Fasc. = inferior longitudinal fasciculus; Mid. Long. Fasc. = middle longitudinal fasciculus; Par. Asl. Tr. = parietal aslant tract; Reticulospi. Tr. = reticulospinal tract; Sup. Long. Fasc. = superior longitudinal fasciculus; Unc. Fasc. = uncinate fasciculus; Vert. Occ. Fasc. = vertical occipital fasciculus

### Myelin water fraction (MWF)

PGS_GI_ was associated with the tracts’ MWF values in four white matter fiber tracts, of which two exhibited positive effects and two exhibited negative effects (see Figure 4b and supplementary Table S3). Higher PGS_GI_ was linked to higher MWF values of the corticopontine tract occipital in the left hemisphere and the reticulospinal tract in the right hemisphere, while higher PGS_GI_ went along with lower MWF values of the cingulum parahippocampal in the left hemisphere and the parietal aslant tract in the right hemisphere. Higher MWF indicates higher myelin content and thus greater myelination. As for ODI, the multiple mediator analysis revealed no white matter fiber tracts that showed a significant association between the tracts’ MWF values and general intelligence (path *b*), and thus no mediators (path *a*\**b*) for the relation between PGS_GI_ and *g* could be identified.

## Discussion

The relation between FA values and intelligence has often been demonstrated, but it is still unclear whether it is due to greater axon density, parallel, homogenous fiber orientation distributions, or greater myelination. Using NODDI and MWF imaging data, we addressed this question and analyzed the microstructural architecture of intelligence in more detail. Furthermore, we were interested whether white matter microstructure indices are involved in the biological pathway that links genetic disposition to phenotype. Thus, we conducted for the first time mediation analyses in which we tested whether NDI, ODI, and MWF of 64 white matter fiber tracts mediated the effects of PGS_GI_ on general intelligence in a large sample of at least 500 healthy young adults. By doing so, we showed that NDI, but not ODI or MWF of white matter fiber tracts was significantly associated with general intelligence and that the NDI of six fiber tracts mediated the relation between genes and *g*.

With regard to the three hypotheses formulated by Stammen et al. [34] on possible links between brain characteristics and intelligence, our results provide clear evidence that differences in neurite density are crucial for differences in intelligence in the white matter, but not differences in neurite orientation dispersion or myelination. Since white matter mainly consists of myelinated axons [147], this means that the number or density of axons is more important for intelligent performance than their arrangement or degree of myelination. We found that 18 white matter fiber tracts showed a significant association between NDI and intelligence. For most of the fiber tracts, the correlation was positive, which fits the hypothesized explanation of Stammen et al. [34] that higher neurite density enables more parallel information processing by providing more possible pathways to think through problems simultaneously. It is a well-established finding that bigger brains are associated with higher levels of intelligence [26,148,149] and a common explanation for this phenomenon is that individuals with more cortical volume are likely to have a higher number of neurons [150]. Since most neurons have only a single axon [151], a higher axonal packing density suggests that there are more neurons in more intelligent individuals, which in turn provide them with enhanced computational power for problem solving and logical reasoning.

Among the white matter fiber tracts that showed a positive association between NDI and *g* were many fiber tracts whose FA values had already been associated with intelligence [26,15,34], namely the cingulum, uncinate fasciculus, superior longitudinal fasciculus, and fornix. This means that our results complement and specify the previous FA results in terms of neurite density. All four fiber tracts have been associated with higher cognitive functions relevant to intelligence, such as attention, cognitive control, memory, visual-spatial functions, or language [152,141,153]. We found significant positive associations between NDI and general intelligence in bilateral parahippocampal parietal, bilateral parolfactory, and left-hemispheric frontal parietal subcomponents of the cingulum. Previous studies have shown that higher NDI in the (para)hippocampal cingulum was positively associated with higher cognition in a mixed sample of healthy older adults and patients with mild cognitive impairment or dementia [154], as well as with episodic memory and processing speed in healthy older adults [54]. The analysis of Raghavan et al. [154] also revealed that the superior longitudinal fasciculus had the strongest positive correlation between NDI and global cognition. This is consistent with our results of positive associations in the superior longitudinal fasciculus 1 in the left hemisphere and in the superior longitudinal fasciculus 3 in the right hemisphere. In contrast, our finding of an association between NDI and the left-hemispheric fornix was not found by Raghavan et al. [154], who averaged the values of the left and right fornix for their analysis. In addition, Coad et al. [53], who analyzed the pre- and postcommissural fornix separately, did not report any association between NDI and various cognitive factors. However, since the existing studies used different cognitive measures, defined the areas of white matter differently, and included different age groups, there are many possible factors that could have caused the different results.

Not previously noticed in FA studies on intelligence were the positive associations with the middle longitudinal fasciculus, the corticopontine tract parietal, and the reticulospinal tract in the left hemisphere. However, they were also often not included as investigated fiber tracts in previous analyses. The middle longitudinal fasciculus is an association fiber tract that connects the superior temporal gyrus with the superior parietal lobule and parietooccipital region and appears to be involved in auditory comprehension as a part of the dorsal auditory stream [141] and higher-order functions related to acoustic information [155]. The corticopontine tract runs from the cerebral cortex through the internal capsule and ends at the unilateral pontine nucleus [156]. It is part of the cerebrocerebellar system and represents an intermediate step in involving the cerebellum into the distributed neural circuits relevant to motor control, thought, and emotion [157]. The corticoreticulospinal tract originates from the premotor cortex, descends to the spinal cord and is part of the extrapyramidal system [158,159]. Although this tract is primarily associated with gross motor function, gait function, and postural stability [159,158], it may also be important for cognitive function in as yet unknown ways, as there is evidence that poor motor function is associated with accelerated cognitive decline in old age [160,161].

However, there were not only tracts whose NDI was positively associated with general intelligence, but also tracts that showed a negative relation, namely the bilateral frontal aslant tract, the left-hemispheric superior longitudinal fasciculus 2, the right-hemispheric thalamic radiation posterior, the left-hemispheric corticostriatal tract superior, and the left-hemispheric vertical occipital fasciculus. The finding that there were both positive and negative associations between white matter fiber tracts’ NDI and intelligence may explain why Genç et al. [52] as well as James et al. [162] found no associations between cognition and total white matter NDI. Although negative associations are less straightforward to explain, we are not the first to have revealed negative associations in circumscribed areas of the brain for structural properties that are generally positively associated with intelligence [72,74]. Furthermore, negative associations between neurite density of fiber tracts and specific skills such as single-word reading and phonological processing have already been reported for children [163]. At first, it seems counterintuitive that lower axonal density was associated with higher intelligence as it contradicts the typical finding that “bigger is better” [26,148,149]. However, it could be the result of efficient wiring and thus signal transmission of the brain so that an optimal balance between signal and noise can be achieved and no redundant or irrelevant information is passed on, which could make efficient problem solving difficult due to inefficient circuitry [52,164]. Establishing and refining efficient brain circuits may be due to regressive events such as axon pruning or synapse elimination [165], which would fit well with the neural efficiency hypothesis stating that higher intelligent individuals need to invest less cortical resources while performing cognitive tasks [166,167]. The vertical occipital fasciculus, for example, is an association fiber tract that runs vertically at the posterolateral corner of the brain and interconnects the dorsal and ventral visual stream [168]. It may be that more precise information exchange between its target regions via less dense axons is advantageous for intelligent thinking, especially since Genç et al. [72] were able to show that lower nodal efficiency and thus less efficient information exchange of the ventromedial visual area 1 with the rest of the brain is associated with higher intelligence. However, it remains to be seen whether future research efforts will be able to replicate our findings of negative associations between NDI of the mentioned fiber tracts and general intelligence in independent samples.

Our result that the orientation dispersion of no fiber tract was related to intelligence is consistent with the results of Callow et al. [55], who used the ICBM-DTI-81 white matter atlas [169–171] and reported a negative association in healthy adults only for the cerebellar peduncle, which was not included in our analysis. Our lack of an association at the level of individual fiber tracts is also well in line with the findings of Genç et al. [52] and James et al. [162], who reported no association at the global level. In contrast, Raghavan et al. [154] showed both significant positive and negative relations between ODI and global cognition in different fiber tracts, but used a mixed sample of healthy subjects and patients for their analysis.

Although it is a long-standing hypothesis that differences in myelination may underlie differences in intelligence [172], our results showed that this is not the case for general intelligence. MWF was not significantly associated with *g* in any fiber tract of the white matter. Penke et al. [20], who first demonstrated an association between myelination in terms of MTR and general intelligence, also showed in their study that this association was completely mediated by a general factor of information processing speed. Accordingly, it could be that MWF is more specifically associated with processing speed than with general intelligence, especially since Page et al. [173] could show that both cognitive domains are at least partially separable with only partly overlapping cerebral correlates. Gong et al. [58] reported associations between lower MWF and steeper declines in processing speed in cognitively unimpaired adults and similar findings have been shown in patients with multiple sclerosis [174–176]. Recent results also suggest correlations of myelination with other cognitive abilities such as executive functions [59] or memory performance [60], so that the subfactors below *g* may be more strongly influenced by differences in myelination. Another way in which myelination could influence cognitive processes could be the coordination of different neuronal networks by accelerating action potentials and mediating precise timing and synchronization between neuronal ensembles [177]. However, more research is needed to understand the role of myelin architecture in functional connectivity [177].

Our finding that PGS_GI_ was significantly associated with NDI of 28, ODI of 16, and MWF of four white matter fiber tracts is in accordance with previously reported results demonstrating relations between genetic variants associated with intelligence and various brain measures [72,178,74,75]. Even if not no association was found between genes for intelligence and myelination, the low proportion of fiber tracts with an association and the lack of an association between myelination and intelligence indicate that differences in intelligence may not mainly be due to differences in myelination, as Lee et al. [70] had already assumed. This result also fits well with the fact that gene sets such as myelin sheath or regulation of myelination were not found to be significantly associated with intelligence in the GWAS used to calculate the PGS_GI_ [64]. Although Schmitt et al. [179], who used a different, less sensitive metric for myelination, found similar spatial heritability patterns for myelination and surface area, they reported that myelination, surface area, and cortical thickness were largely genetically independent in adults. Studies revealed that PGS_GI_ was associated with surface area and cortical thickness of various brain areas and that both properties of specific brain regions mediated the association between PGS_GI_ and intelligence [75,72,74]. Thus, it is conceivable that genes relevant to intelligence overlap more with genes relevant to structural morphological brain properties but less with genes relevant to myelination. However, it could also be that genes that overlap with intelligence and myelination have not yet been found in GWAS due to small effect sizes and limited sample sizes.

Interestingly, the neurite density of six fiber tracts mediated the association between PGS_GI_ and general intelligence. Five of them, namely the left-hemispheric middle longitudinal fasciculus, the bilateral cingulum parahippocampal parietal, the left-hemispheric uncinate fasciculus, and the right-hemispheric superior longitudinal fasciculus 3 exhibited positive mediation effects, which are well in line with the results of previous mediation analyses [75,72,74]. The middle longitudinal fasciculus runs from the superior temporal gyrus to the superior parietal lobule and parietooccipital region [141], the cingulum parahippocampal parietal from the medial temporal lobe to the parietal and occipital lobes [180], the uncinate fasciculus from the anterior temporal lobes and amygdala to the lateral orbitofrontal and anterior portion of the prefrontal cortex [141], and the superior longitudinal fasciculus 3 from the frontal and opercular areas to the supramarginal gyrus [141]. Genç et al. [72] identified the bilaterally averaged surface area of the superior medial parietal cortex, intraparietal areas, and the posterior temporal cortex as positive mediators between PGS_GI_ and intelligence and classified all of them as part of the P-FIT model [13]. Furthermore, they found the structural network efficiency of the inferior frontal gyrus to mediate positively between genes and intelligence. Lett et al. [75] reported that the association between genetic variants and general intelligence was mediated positively by the cortical thickness and surface area of the anterior cingulate cortex, the prefrontal cortex, the insula, the medial temporal cortex, and the inferior parietal cortex. Similar mediating areas were found by Williams et al. [74]. Our results of positive mediating white matter fiber tracts thus connect regions of the brain whose morphological properties have already been identified as mediating factors between genes and intelligence and as relevant for intelligent thinking in the P-FIT model [13]. Jung and Haier [13] assumed that the entire process of reasoning depends on the fidelity of underlying white matter and we provided evidence that a higher axonal packing density of specific white matter fiber tracts is one factor that links the genetic basis of intelligence to the corresponding phenotype.

One white matter fiber tract, namely the left-hemispheric frontal aslant tract had a negative mediation effect, which was due to a negative association between its NDI and general intelligence. Negative mediation effects were also reported by Genç et al. [72], who, for example, identified the surface area of the inferior frontal sulcus as a negative mediator between genetic variants associated with educational attainment and intelligence. The frontal aslant tract runs from the pars opercularis and pars triangularis of the inferior frontal gyrus and the anterior insula to the supplementary motor area (SMA) and pre-SMA and is believed to be involved in speech planning, initiation, and production, but also kinematics and visuomotor processes [141]. As intraoperative direct electrical stimulation of the left frontal aslant tract led to stuttering [181], it could be that precise and efficient information exchange between its target regions due to lower axonal density is crucial for optimal functioning. However, the frontal aslant tract has not yet been well studied and has only recently been linked to other cognitive functions, so its role in cognition needs to be further characterized [182].

There are certain limitations to our study. Our paper is limited in its population representativeness as our sample mainly consisted of German university students who had a mean IQ score one standard deviation above average which might have impacted the associations between genetic variants, the brain, and intelligence we observed. As neurite density as well as myelin content are associated with age [141,183], future studies should examine whether our results, which were limited to young adulthood, expand to other age groups or even use longitudinal designs. Furthermore, our analyses were restrained to individuals of European ancestry, thus further research is needed to investigate whether the NDI of the same white matter fiber tracts mediates the association between PGS_GI_ and general intelligence to the same degree across ancestries. As PGS_GI_ only predicted up to 5.2% of variance in general intelligence in independent samples [64], it could also be that additional white matter fiber tracts will be found when using a PGS that is based on a larger sample size and has better predictive power for intelligence. Additionally, our measures of neurite density, neurite orientation dispersion, and myelination were based on neuroimaging techniques that are limited in spatial resolution. It is known that the human brain contains different classes of axons ranging from large-diameter myelinated to small-diameter unmyelinated fibers [184,185], which could not be differentiated by the methods used. Finally, we limited our study to analyzing the relation between PGS_GI_, white matter microstructural architecture, and general intelligence in healthy subjects. However, our type of analysis can be extended to the multitude of other PGS [186], brain correlates, and phenotypes (e.g., intelligence subfactors) in future studies and may also provide valuable insights for clinical samples.

The present paper provides the first study examining the mediating effects of the white matter microstructural indices NDI, ODI, and MWF on the association between genetic variation and general intelligence. We showed that the neurite density of specific white matter fiber tracts played a mediating role in the relation between cumulative genetic load for general intelligence and *g*-factor performance. In contrast, we found no significant associations with general intelligence for ODI and MWF. The latter was surprising, as myelination was considered a possible neurobiological correlate of intelligence. These findings are a crucial step forward in decoding the neurogenetic underpinnings of general intelligence, as they identify that the neurite density of specific white matter fiber tracts relate polygenic variation to *g*.

## Acknowledgements

The authors would like to thank Wendy Johnson for providing the *g* factor scores, all research assistants for their support during the behavioral measurements, PHILIPS Germany (Burkhard Mädler) for the scientific support with the MRI measurements as well as Tobias Otto for technical assistance.

## Statements and Declarations

### Author contribution

E.G., R.K., and O.G. conceived the project and supervised the experiments. E.G., C.St., R.K., and O.G. designed the project. R.K. and J.S.P. planned and performed genetic experiments. C.Sc. and C.F. collected data. C.St., E.G., J.S.P., D.M., and M.J.H. analyzed the data. C.St. and E.G. wrote the paper. All authors discussed the results and edited the manuscript.

### Data availability statement

The data that support the findings of this study are available from the corresponding author upon reasonable request or can be downloaded from an Open Science Framework repository [https://osf.io/29cp6/]. We will release the data after our manuscript has been accepted for publication.

### Ethics approval

The study was performed in line with the principles of the Declaration of Helsinki. Approval was granted by the Ethics Committee of the Faculty of Psychology at Ruhr University Bochum (vote Nr. 165).

### Consent to participate

Informed consent was obtained from all individual participants included in the study.

### Consent to publish

The authors affirm that human research participants provided informed consent for publication of the images in Figure 2.

