## Supplemental Figure 1 for "Neurite density but not myelination of specific fiber tracts links polygenic scores to general intelligence"

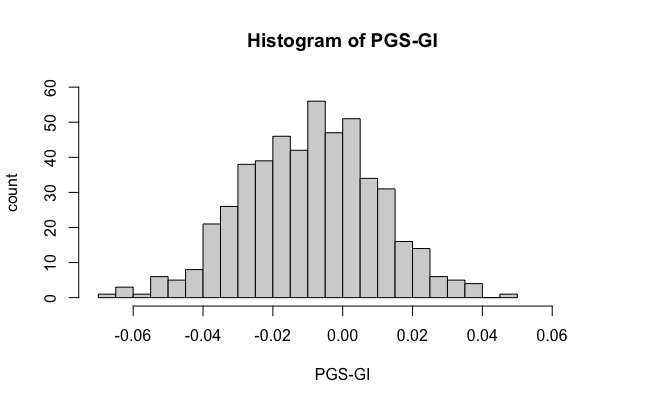
**Supplementary Fig. 1** Distribution of the polygenic score for general intelligence (PGS_GI_) in our sample
